# Early steps of embryo implantation are regulated by exchange of extracellular vesicles between the embryo and the endometrium

**DOI:** 10.1101/2022.04.28.489970

**Authors:** Joanna Szuszkiewicz, Kamil Myszczynski, Zaneta P Reliszko, Yael Heifetz, Monika M Kaczmarek

## Abstract

In early pregnancy, as the embryo arrives in the uterus, intensive communication between the embryo and uterus begins. Hundreds of molecules are known to be involved, but despite numerous findings, full understanding of the complexity of the embryo–maternal dialog remains elusive. Recently, extracellular vesicles, nanoparticles able to transfer functionally active cargo between cells, have emerged as important players in cell–cell communication, and as such, they have gained great attention over the past decade also in reproductive biology. Here we use a domestic animal model (*Sus scrofa*) with an epitheliochorial, superficial type of placentation because of its advantage in studding uterine luminal fluid extracellular vesicles. We show that during early pregnancy, the uterine lumen is abundant with extracellular vesicles that carry a plethora of miRNAs able to target genes involved in embryonic and organismal development. These extracellular vesicles, upon the delivery to primary trophoblast cells, affect genes governing development as well as cell-to-cell signaling and interactions, consequently having an impact on trophoblast cell proliferation, migration, and invasion. We conclude that exchange of a unique population of extracellular vesicles and their molecular cargo at the maternal–embryo interface is the key to the success of embryo implantation and pregnancy.

## Introduction

The maternal recognition of pregnancy, followed by embryo implantation, is a crucial step in securing a pregnancy to full term in mammals. Our current understanding of such processes is mostly rooted in in vitro models of human cells and animal studies, with livestock gaining more attention in providing important insights into embryo–maternal interaction (1). Acquisition of the uterine receptivity and timely development of an embryo is necessary for the establishment of synchronized molecular communication at the embryo–maternal interface, understood as a precise spatiotemporal exchange of a broad range of biologically active compounds synthesized and secreted by the endometrium and/or embryo (2, 3). Importantly, at this heavily complex and yet not fully understood stage of pregnancy, high mortality of the embryo occurs. In most mammals, including swine, sheep, cattle, and humans, from 10% to 40% of pregnancies fail due to the losses occurring during the peri-implantation period of pregnancy (4–8); however, still much remains to be discovered to broaden our knowledge of the complexities of early pregnancy events.

Although implantation strategies differ among eutherian mammals, initial stages of apposition and adhesion are common. Rodents’ and primates’ embryos almost immediately attach to receptive epithelium after entering the uterine cavity. On the other hand, domestic animals have a protracted pre-implantation period, characterized by the establishment of an intimate dialog between the embryo and the uterus (9). In pigs, during the peri-implantation period, embryos migrate within the uterus before apposition, grow and undergo a dynamic morphological transformation, which all demand an extensive regulation of cell behavior (e.g., differentiation, proliferation, and migration) (9–11). At the same time, extensive changes within the uterine walls take place to reach full receptivity and enable successful attachment and implantation of developing embryos. As in other mammals, in pigs, the expression of a number of endometrial genes is altered during the peri-implantation period, having its consequence in time- and space-dependent modifications of surface molecules and secretion of others (12, 13). This is an example of a highly coordinated spatiotemporal intercellular communication, providing a nurturing environment for establishment and maintenance of pregnancy, which requires extensive intercellular communication.

Among the wide range of cell–cell communication strategies, there is one, relatively recently discovered, governed by extracellular vesicles (EVs). EVs are the heterogeneous in size, origin, and content population of nano-sized cell-derived membrane vesicles (14). EVs are able to transfer, between even distant cells, their biological cargo consisting of proteins, lipids, and nucleic acids (15). They have emerged as important players in numerous pato- and physiological processes, and as such, they have gained great attention over the past decade (16). EVs have already gained attention among reproductive biologists, providing increasing data on their specific roles (17–19). To date, EVs have been identified as being released by embryos from the earliest days after fertilization (20–22), through the peri-implantation phase (23–27), to the stages when placenta is already formed (28). EVs have also been studied in various reproductive pathophysiological conditions and diseases, including preeclampsia (29).

The cargo of EVs is crucial for affecting recipient cells (30, 31). EVs are known transporters of a variety of molecules, including nucleic acids, proteins, and lipids. Although the term exosomes (one of the EV subtypes) has been known since the 1980s (32), the real breakthrough in the field of EVs happened later, after the discovery that EVs are able to transport functional mRNA and microRNAs (miRNAs) and affect the function and fate of neighboring and distant cells (33). miRNAs are short, noncoding regulatory RNAs able to affect gene expression (34). To date, a great number of miRNAs have been identified as important regulatory molecules during pregnancy in mammals, including pigs (35). Previously, we identified embryonic (23) and endometrial (36) miRNAs showing a potential to affect genes not only in the place of synthesis but also in distant cells because of the ability of miRNAs to travel via circulation (37, 38). The potential roles of miRNAs transported by EVs in maternal–embryo communication have been already studied in several mammals (20, 24, 27, 39, 40). Nevertheless, many of the presented reports lack systematic characterization of EVs and identification of their functions during the peri-implantation phase, when common stages of embryo apposition and adhesion occur in mammals. In addition, the composition of miRNA cargo secreted by both sites of the early dialog (i.e., the embryo and endometrium) has not yet been reported.

To characterize EVs present in the uterus during peri-implantation period and define their impact on trophoblast cells physiology upon *in vitro* autotransplantation, we used a porcine model with an epitheliochorial, superficial type of placentation. Because the pig has the most superficial placenta and lacks significant invasion of the uterine luminal epithelia, we could follow the early cell-to-cell communication between the embryo and the endometrium. Here, for the first time, we show that the uterine cavity during early pregnancy is abundant in EVs containing numerous miRNAs that can be internalized by porcine trophoblast cells, wherein they affect gene expression, stimulate proliferation, and inhibit migration and invasion. These studies further support our hypothesis that processes governing early steps of embryo implantation are regulated via molecular cargo carried by EVs and exchanged at the embryo– maternal interface, being a critical element of an intimate crosstalk between the embryo and the uterus.

## Materials and Methods

### Material

Crossbred gilts (Pietrain × Duroc) of similar age (8 – 9 months) and genetic background from one commercial herd were artificially inseminated 12 h (Day 0) and 24 h after the first signs of the second estrus. Samples were collected in the slaughterhouse on Days (D) 12, 14, and 16 of pregnancy. The day of pregnancy was confirmed by the size and morphology of conceptuses, as described previously (23, 36). Each horn of the uterus was flushed twice with 20 ml of 0.01 M phosphate-buffered saline (PBS; pH 7.4). Uterine luminal flushings (ULFs) were collected and immediately transported to the laboratory on ice for EVs isolation. Fragments of uterus and embryos were cut and placed immediately in 4% paraformaldehyde solution for immunofluorescence staining. Conceptuses from D16 were transferred into Dulbecco’s Modified Eagle Medium/Nutrient Mixture F-12 Ham (DMEM/F12; Sigma-Aldrich) medium supplemented with 1% (v/v) penicillin/streptomycin (P/S; Sigma-Aldrich) and immediately transported to the laboratory.

All procedures involving animals were conducted in accordance with the national guidelines for agricultural animal care in compliance with EU Directive 2010/63/UE.

### Immunofluorescence

Paraformaldehyde-fixed and paraffin-embedded tissues were sectioned (4 μm) and mounted on chromogelatin-precoated slides (Menzel-Glaser). Sections were deparaffinized, rehydrated, and blocked using 10% normal donkey serum (Jackson Immunoresearch). Next, slides were incubated with primary mouse monoclonal anti-CD63 antibody (Abcam, cat. # ab8219, 1:30) overnight at 4 °C. Secondary antibody, Cy3-conjugated donkey anti-mouse IgG (Jackson ImmunoResearch, cat. # 715-165-150, 1:1 000) was applied and incubated for 1 hr at room temperature. Negative controls were performed without primary antibodies. Finally, sections were mounted in Ultra Cruz Mounting Medium with DAPI (Santa Cruz Biotechnology) and visualized with an epifluorescent microscope Zeiss Axio Imager System (Carl Zeiss).

### Extracellular vesicles isolation

After embryo decantation, ULFs were collected and subjected to stepwise centrifugation at 4°C (156 x g, 10 min; 2 000 x g, 10 min; 10 000 x g, 30 min) to eliminate dead cells and cell debris. Final supernatants were passed through 0.22 µm filters. 40 ml of filtrates were ultracentrifuged twice in sterile 10.4 mL polycarbonate bottles at 100 000 x g for 70 min at 4°C using an Optima L-100 XP Ultracentrifuge equipped with 90 Ti Fixed-Angle Titanium Rotor (Beckman Coulter). The final EVs pellets were suspended in 200 µl of PBS (Lonza). Samples were aliquoted and stored at -80°C for further analysis.

### Nanoparticle Tracking Analysis

Nanoparticle Tracking Analysis (NTA) was performed by a NanoSight NS300 (NanoSight Ltd) equipped with a 405 nm laser and an automatic syringe pump system. Samples were diluted in PBS (Lonza) to reach a particle concentration suitable for unbiased analysis. Three 60 sec videos were recorded of each sample with camera level 13 and the detection threshold set at 3. Videos were analyzed with NTA software version 3.4 to determine the concentration and size of measured particles with corresponding standard error. Particle concentration and mode of particle size in each sample were used for the statistic. The number of biological replicates was 4-5 per group, as indicated in the graph. Differences between groups were tested with ordinary one-way ANOVA and Tukey’s multiple comparisons test.

### Transmission electron microscopy

Isolated EVs were diluted with PBS (Lonza) and loaded onto formvar-carbon–coated copper grids. Samples were stained with 1% uranyl acetate for 1-2 min and dried at room temperature. Images were obtained using a Tecnai 12 transmission electron microscopy (FEI), operating at an acceleration voltage of 100 kV, equipped with a CCD camera MegaView II. Separate images were taken to provide wide-field images, showing the whole population of vesicles or close-up images of a single vesicle. Three samples per group were visualized.

### Western blotting

Protein concentrations in EVs samples were determined using the Bradford assay. The volume of EVs containing 30-40 µg of proteins was mixed with RIPA buffer (PBS pH 7.4, 1% Triton X-100, 0.5% sodium deoxycholate, 0.1% sodium dodecylsulfate, 1 mM EDTA) and incubated 15 min at 4°C. After adding Loading Buffer (BioRad), samples were boiled at 95°C for 5 min. Proteins were separated by SDS-PAGE (SDS-polyacrylamide gel electrophoresis) and transferred to PVDF membranes and blocked in 5% non-fat dry milk in TBS-T (Tris-buffered saline, containing 0.1% Tween-20). The membranes were incubated with the following antibodies (Abcam): mouse monoclonal anti-CD63 [MEM-259] (cat. # ab8219, 1:170), mouse monoclonal anti-HSP70 [3A3] (cat. # ab5439, 1:500), rabbit polyclonal anti-Syntenin (cat. # ab154940, 1:500), mouse monoclonal anti-TSG101 [4A10] (cat. # ab83, 1:100) as positive EVs markers or rabbit polyclonal anti-calreticulin (cat. # ab15607, 1:200), rabbit polyclonal anti-AGO2 (cat. # ab32381, 1:500) as negative EVs markers. Signal was visualized with Clarity Western ECL Substrate and ChemiDoc Imaging System (Bio-Rad).

### Porcine trophoblast primary cells isolation

Porcine trophoblast primary cells (pTr) were isolated according to the method established previously (41). Briefly, conceptuses were washed three times in DMEM/F12 medium and digested with 0.25% trypsin (Biomed) for 30 min. The suspension was filtered through sieve mesh and centrifuged 200 x g for 10 min at 8°C. After washing, the cell pellet was resuspended in DMEM/F12 and the cell number was estimated with trypan blue staining (mean % of dead cells ± SEM = 9.07 ± 0.42). Cells were cultured in DMEM/F12 supplemented with 10 % (v/v) newborn calf serum (NCS; Sigma-Aldrich) and 1 % (v/v) P/S at 37°C with 5% CO_2_.

### Scratch wound healing assay

To test the effect of EVs on basic pTr cell migration a scratch wound healing assay was used in in vitro autotransplantation approach, meaning that EVs and trophoblast cells were isolated from the same animal (n=3). First, 2 × 10^5^ cells per well were seeded on 24 well plates (collagen I-coated plates; Corning BioCoat) and cultured to reach about 70-80% confluence. For treatment, the medium was replaced by free DMEM/F12 supplemented with 0.2, 2, or 4 % of EVs (v/v) for 6 or 12 h. After this time, wounds were struck through confluent cell monolayers using a pipette tip. Next, cells were rinsed with PBS and placed in DMEM/F12 supplemented with 10 % (v/v) NCS and 1 % (v/v) P/S. To observe wound healing, pictures were taken at 0 h and every 3 h within a 24 h-long time frame using the Axio Observer and ZEN 2.5 blue (Zeiss). The average distance between the two margins of the scratch was measured using ImageJ (42). The motility ratio was calculated at each time point according to the following formula: distance at a specific time point - distance at 0 h. The area under the curve was analyzed using a repeated measures one-way ANOVA with Dunnett’s multiple comparisons test (vs. control). The assay was repeated three times independently.

### Transwell invasion assay

Transwell invasion assay was performed as described elsewhere (43) with slight modifications. Briefly, 1.5 × 10^5^ pTr cells were seeded in the top chamber of 8 μm pore size Matrigel-coated inserts (Corning). After reaching the confluence, cells were treated with 2% EVs for 6 h in DMEM/F12. As for wound healing assay, each treatment was performed using EVs and trophoblast cells isolated from the same animal (n=6). In the lower chamber, 20% NCS in DMEM/F12 served as a chemoattractant. Negative controls were performed with either no EVs or no chemoattractant added. The cells were allowed to migrate for 18 h, and then, the cells located on the membrane in the lower chamber were fixed in 4% paraformaldehyde for 10 min, stained with Hoechst33342 (2 µg/ml; Thermo Fisher), visualized using Axio Observer and ZEN 2.5 blue (Zeiss) and counted manually. Repeated measures one-way ANOVA and Dunnett’s multiple comparisons test (vs. negative control) were used to find statistical significance.

### Proliferation assay

CellTiter 96 AQueous One Solution Reagent (Promega) was used to assess the effect of EVs on cell proliferation rate in in vitro autotransplantation approach (n=4). The pTr cells were treated with 2% EVs for 6 h in DMEM/F12 and medium was changed to DMEM/F12 supplemented with 10 % (v/v) NCS and 1 % (v/v) P/S. After 18 h of additional culture, CellTiter was added, and absorbance was measured at 490 nm. The cells treated with 20% NCS were used as a positive control. Repeated measures one-way ANOVA and Dunnett’s multiple comparisons tests (vs. control) were used to find statistical significance. The proliferation assay was performed in quintuplicates and was repeated five times independently.

### Gene expression analysis

The pTr cells were treated for 6 h with 2% EVs in free DMEM/F12 and next cultured in DMEM/F12 supplemented with 10 % (v/v) NCS and 1 % (v/v) P/S for another 18 h on 6-well plates (collagen I-coated plates; Corning BioCoat). As for other in vitro tests, each treatment was performed using EVs and trophoblast cells isolated from the same animal (n=5). After treatment, total RNA was isolated using miRVana miRNA isolation kit. Next, its quantity and quality were assessed using Bioanalyzer 2100 (Agilent Technologies) and Qubit 3.0 Fluorometer (Thermo Fisher Scientific). For RNA-seq, one microgram of total RNA was used to construct cDNA libraries with the TruSeq Stranded Total RNA with Ribo-Zero Gold Human/Mouse/Rat kit (Illumina). Sequencing was performed using Illumina NovaSeq 6000 (coverage - 50 x, read length - 150 bp PE = paired-end; Macrogen Europe BV). The raw reads were first quality checked and cleaned up using FastQC (44) and Trimmomatic (45) to obtain high-quality reads. The reads were mapped to the *Sus scrofa* reference genome using STAR (46). Afterward, the mapped reads were counted with featureCounts 2.0.3 (46) and differential gene expression analysis was conducted using DESeq2 1.34.0 (47). The absolute value of fold change ≥ 1.27 and adjusted p-value < 0.05 were used as a criterion to identify differentially expressed genes. The results of the analysis were visualized using R (48).

RNA-seq results were validated using TaqMan Gene Expression Assays (see Supplementary Table 1A) and TaqMan One-Step RT-PCR Master Mix Reagents Kit (Life Technologies). The reaction was performed using 10 ng of total RNA as a template. Negative controls without a template were performed in each run. RT-qPCR reactions were performed in duplicates on the ABI HT7900 real-time PCR system (Life Technologies). The expression values were calculated including the efficiency of the reactions (49) and normalized to the reference gene (*GAPDH*) showing the best stability (0.032), calculated by NormFinder (50). Person correlation of RNA-seq and RT-qPCR data using a log2 mean-fold change was calculated.

### miRNA cargo characterization

Total RNA enriched in small RNA fraction was isolated from EVs prepared as described above using mirVana miRNA Isolation Kit (Life Technologies). The quantity, quality, and size distribution of total and small RNAs were assessed using Bioanalyzer 2100, and RNA 6000 Pico and Small RNA kits (Agilent Technologies). Custom-designed TaqMan Low Density Arrays (TLDA, Life Technologies) (38) were used to profile miRNA cargo carried by EVs. Two-card sets were used, which contained 11 reference and 166 investigated mature miRNAs, selected based on our previous studies and literature (see Supplementary Table 2). RT-PCR reactions were performed according to the manufacturer’s instructions. Briefly, 21 ng RNA per reaction was reversely transcribed with TaqMan miRNA Reverse Transcription Kit (Life Technologies) and custom RT Primer Pool. Next, RT product mixed with TaqMan Universal Master Mix II (no AmpErase UNG) was loaded into array card. ViiA™ 7 Real-Time PCR System (Life Technologies) was used to perform RT-qPCR reaction. Raw fluorescent data, imported from SDS 2.4 software were analyzed with PCR Miner (49) to calculate reaction efficiency. Ct values□L35 were considered to be below the detection level and were excluded from further analysis, only if Ct values were consistently low in all days tested. The most stable reference miRNA was chosen using NormFinder (50) among list of five candidates (ssc-miR-20a-5p, cgr-miR-140-3p, ssc-miR-140-3p, ssc-miR-16, and U6-snRNA). Relative miRNAs expression was normalized using ssc-miR-140-3p, showing the best stability values (card 1 = 0.173; card 2 = 0.030). Values multiplied by 100 were log2 transformed and used to create a circular heatmap of all detected miRNAs (51). For those miRNAs, which expression was not detected (Ct value □L35) on the particular day tested; values were set as 0 and not used for further statistics.

TLDA results were validated using TaqMan MicroRNA Assays (Life Technologies). Briefly, 10 ng of RNA was reversely transcribed with TaqMan MicroRNA Reverse Transcription Kit containing Multiscribe RT enzyme and appropriate primers for each miRNA (see Supplementary Table 1B). For each RT-qPCR reaction, 2.6 µl of cDNA was used along with TaqMan Universal Master Mix II and miRNA probes. Negative controls were performed in each run, without template or reverse transcriptase added. RT-qPCR reactions were performed in duplicates on the ABI HT7900 real-time PCR system (Life Technologies). The expression values were calculated including the efficiency of the reactions (49) and normalized to reference gene ssc-miR-140-3p.

Statistical analysis for TLDA and RT-qPCR was performed using either ordinary two-way ANOVA followed by a Tukey’s multiple comparisons test (for D12, D14, and D16 comparison) or t-test (for D14 and D16 comparison).

### Functional enrichment analysis

A lists of miRNAs detected in EVs and differentially expressed mRNAs after treatment of pTr cells with EVs, along with their fold change (log2) and significance values were uploaded to Ingenuity Pathway Analysis (IPA; Qiagen) tool. In order to increase the clarity of the results, enrichment analysis was performed with excluded chemicals and biological drug interactions and using only experimentally observed interactions. Fisher’s Exact test was used to assess p-values for enriched functions (threshold P < 0.05). Molecule Activity Predictor tool within IPA was applied for downstream effects prediction of differentially expressed molecules after pTr cells treatment with EVs.

### *In silico* miRNA-mRNA interaction analysis

In order to generate list of genes differentially expressed in conceptuses at D15 vs. D12, the raw reads produced by Zhang et al (52), deposited in the NCBI Sequence Read Archive (accession number is PRJNA646603), were first quality checked and cleaned up using FastQC (44) and Trimmomatic (45) to obtain high-quality reads. The reads were mapped to the *Sus scrofa* reference genome using STAR (46). Afterward, the mapped reads were counted with featureCounts 2.0.3 (46) and differential gene expression analysis was conducted using DESeq2 1.34.0 (47). The absolute value of log2 fold change ≥ 1 and adjusted p-value < 0.05 were used as a criterion to identify differentially expressed genes.

For miRNA-mRNA interactions, miRWalk database (53) was utilized. Names of all detected EVs miRNAs were translated into human equivalents, if possible, based on 100% sequence similarity. Out of 79 detected EVs miRNAs, 60 were recognized and used for analysis. From the list of possible miRNA-mRNA interactions, downregulated genes in conceptuses at D15 *vs*. D12 were subtracted. Cytoscape 3.9.1 (54) was applied to visualize potential miRNA-mRNA interactions. Additionally, a list of possible miRNA-mRNA interactions was screened for the presence of genes downregulated in pTr cells after in vitro incubation with EVs and uploaded to create a network in Cytoscape 3.9.1. For subtracting interesting clusters, ClusterViz (55) with the EAGLE algorithm was applied.

### Confocal microscopy

The pTr cells were seeded on cell imaging cover glasses with four chambers (Eppendorf; 10^5^ cells/chamber). RNA cargo carried by EVs prepared as described above was stained with thiazole orange as described elsewhere, with minor modifications (56). Briefly, 10 µl of EVs was incubated with 0.2 µg stain in PBS at 37°C for 30□min. Labeled vesicles were then washed in PBS and ultracentrifuged at 100 000 x g for 70 min (SW 40 Ti Swinging-Bucket Rotor; Optima L-100 XP Ultracentrifuge; Beckman Coulter). Next, the vesicle pellet was resuspended in PBS, added to the pTr cells, and incubated for 3 h at 37°C with 5% CO_2_. Afterward, nuclei were stained with Hoechst 33342 (2 µg/ml; Thermo Fisher) for 30 min, and slides were further incubated with 0.25% Triton X-100 (Sigma-Aldrich) for 10 min. After F-actin staining with Alexa Fluor 488 Phalloidin (5 U/slide; Thermo Fisher) cells were covered with an antifade mounting medium (Vector Laboratories) and placed on a microscopic slide (Menzel-Glaser). EVs RNA cargo uptake was confirmed using LSM800 Airyscan and ZEN 2.5 blue software (Zeiss). Treatment with PBS labeled as described above was used as a negative control.

### Statistical analysis

Statistical analysis, if not stated otherwise, was carried out using GraphPad Prism 8 (GraphPad Inc.). Effects were considered significant at P□<□0.05. Data are presented as mean + standard error of the mean (SEM), except line graphs where mean ± SEM are presented. Sample sizes and other statistical details are indicated in the figures/figure legends.

## Results

### EV populations released to the uterine lumen during peri-implantation period have different characteristics

CD63, a well-known tetraspanin involved in many cellular processes, including vesicular trafficking, is considered a reliable marker of exosomes secretion (30). To determine whether endometrium and conceptuses on D12, D14, and D16 of pregnancy could be the source of EVs in the uterine lumen, we performed histological localization of the CD63 signal. We found that CD63+ cells were present in all examined tissues and days of pregnancy (Fig. 1A). In the endometrium, CD63+ cells were detected in the luminal epithelium. Interestingly, the cellular distribution of the CD63 signal changed from evenly diffused around the whole cell on D12 to polarized to the apical surface and more intense on D16. A similar spatiotemporal pattern was observed in conceptuses, as on D16, the CD63 signal was unevenly distributed, but a stronger signal was observed at the apical site of trophoblast cells. These results indicate that along with establishment of an intimate contact between the luminal epithelium and trophoblast, the extensive trafficking of vesicles takes place at the apical surfaces of cell types.

**Figure 1.**
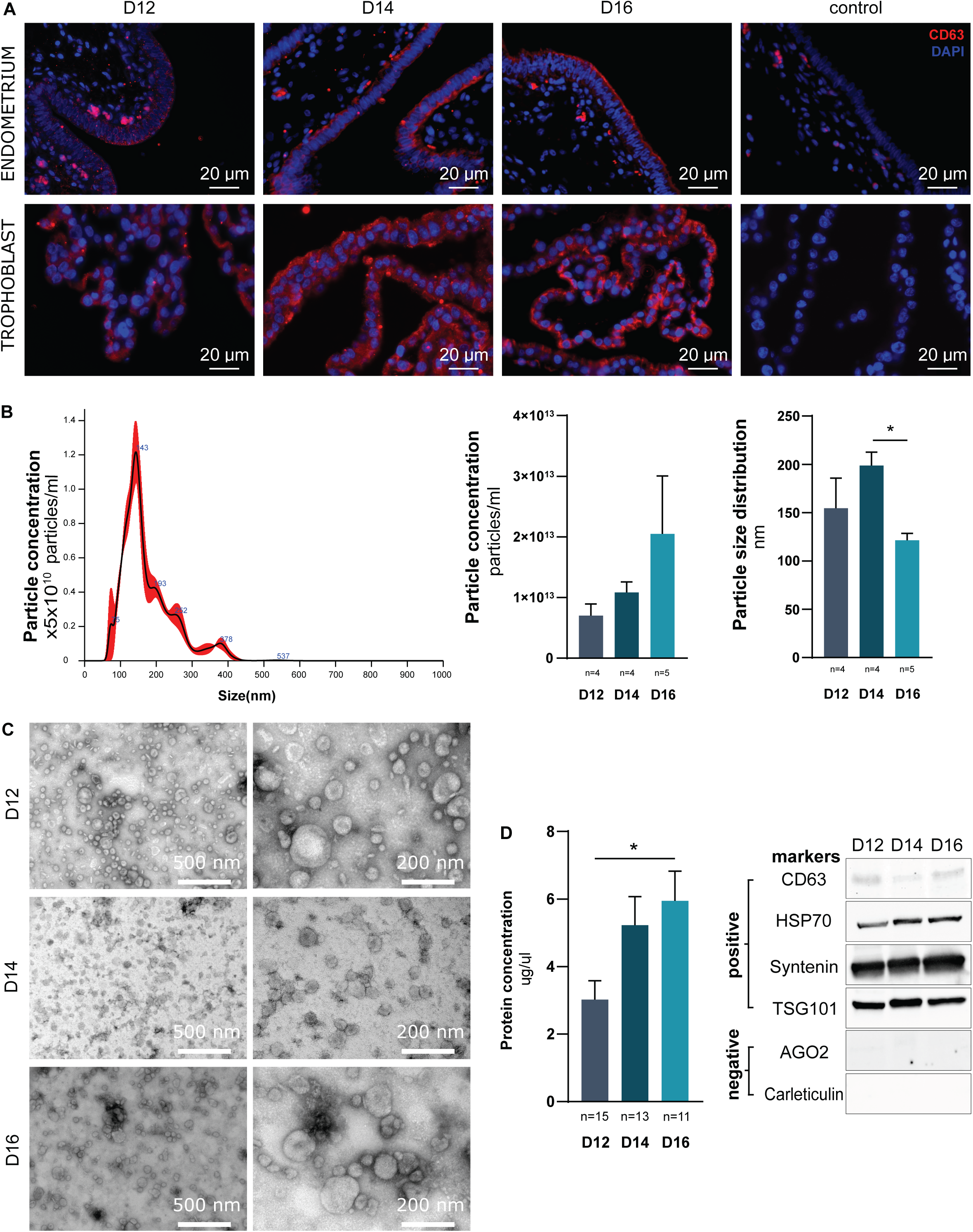
The uterine lumen of early pregnancy is abundant in EVs released by the both endometrium and conceptuses. **A**. Immunolocalization of CD63+ cells in the endometrium (top) and conceptuses (bottom) during consecutive days (D) of pregnancy (D12, D14 and D16). Control staining was performed without primary antibodies. **B**. Nanoparticle Tracking Analysis was used to measure the size and concentration of EVs derived from the porcine uterine lumen at D12, D14, and D16 of pregnancy. Representative profile of particle distribution (left) along with statistical analysis of particle concentration (middle) and particle size distribution (right) for all tested days of pregnancy are presented. * P = 0.0343 (one-way ANOVA and Tukey’s multiple comparisons test). **C**. Transmission electron microscope images for EVs collected from uterine lumen on D12, D14, and D16 of pregnancy. Widefield (left) and close-up (right) images are shown. **D**. The protein concentration for EVs samples collected during pregnancy (D12, D14, D16) supported by the detection of EV protein markers (CD63, HSP70, Syntenin, TSG101). AGO2 and Carleticulin were used as negative markers. * P = 0.0265 (one-way ANOVA and Tukey’s multiple comparisons test).

Once we confirmed that both embryos and the endometrium could be a source of EVs during early pregnancy, we isolated and characterized EVs present in the uterine lumen on D12, D14, and D16. We measured the concentration and size distribution of isolated particles (Fig. 1B). All samples contained a high concentration of nanosized particles, and there were no differences in particle concentration among the analyzed days of pregnancy. Mean particle size was consistent with the size of EVs (D12 = 154.6 nm, D14 = 198.8 nm, D16 = 121.4 nm). Interestingly, the size of D16 particles was significantly smaller when compared with the size of D14 particles (P = 0.0343). To ascertain the morphology of uterine-derived EVs, transmission electron microscopy was performed (Fig. 1C). Images showed an EV population heterogeneous in size with a characteristic artefactual cup shape.

To further characterize the EV population on each day of pregnancy examined, we measured the total protein concentration. We found that the protein level in D16 EVs was significantly higher when compared with D12 (P = 0.0265, Fig. 1D). Moreover, uterine-derived EVs were positive for several proteins—CD63, HSP70, syntenin, and TSG101 (positive EV markers)— and negative for AGO2 and calreticulin (negative EV markers; Fig. 1D).

### Uterine-derived EVs affect trophoblast cells’ migration, invasion, and proliferation

Next, we decided to test EVs’ effect on principal processes governing conceptus growth, spacing in the uterus, and implantation. To accomplish this task, we prepared uterine lumen EVs and pTr cells that originated from the same animal at D16 of pregnancy (in vitro autotransplantation approach). First, we performed a time- and dose-dependent scratch wound-healing experiment. After creating a scratch in the monolayer of pTr cells, we incubated them with a medium supplemented with 0.2, 2, or 4% of EVs for 6 h or 12 h. Media supplemented with 2% of EVs significantly decreased pTr cells’ migration as early as 6 h post treatment (P = 0.0018), whereas a lower EV concentration (0.2%) was effective after 12 h (P = 0.025; Fig. 2A). In subsequent in vitro autotransplantation experiments, we used media supplemented with 2% of EVs in 6-h incubations. We next investigated the invasive capacity of pTr cells under the effect of 20% NCS as a chemoattractant and 2% EV treatment. As expected, 20% NCS induced cell invasion (vs. cells without 20% NCS, P = 0.05; Fig. 2B). To evaluate the effect of EV treatment, we compared the invasion of untreated cells with that of cells incubated with 2% of EVs in the presence of the chemoattractant (20% NCS). Consequently, a significant decrease of invasion rate was indicated when cells were treated with EVs (P = 0.02; Fig. 2B). Further experiments showed that pTr cells incubated with 2% EVs for 6 h exhibited increased proliferation (P = 0.031; Fig. 2C). Taken together, the results indicate EVs present in the uterine lumen of pregnant animals can affect crucial aspects of trophoblast physiology, having an impact on processes governing its development and behavior during initial stages of pregnancy.

**Figure 2.**
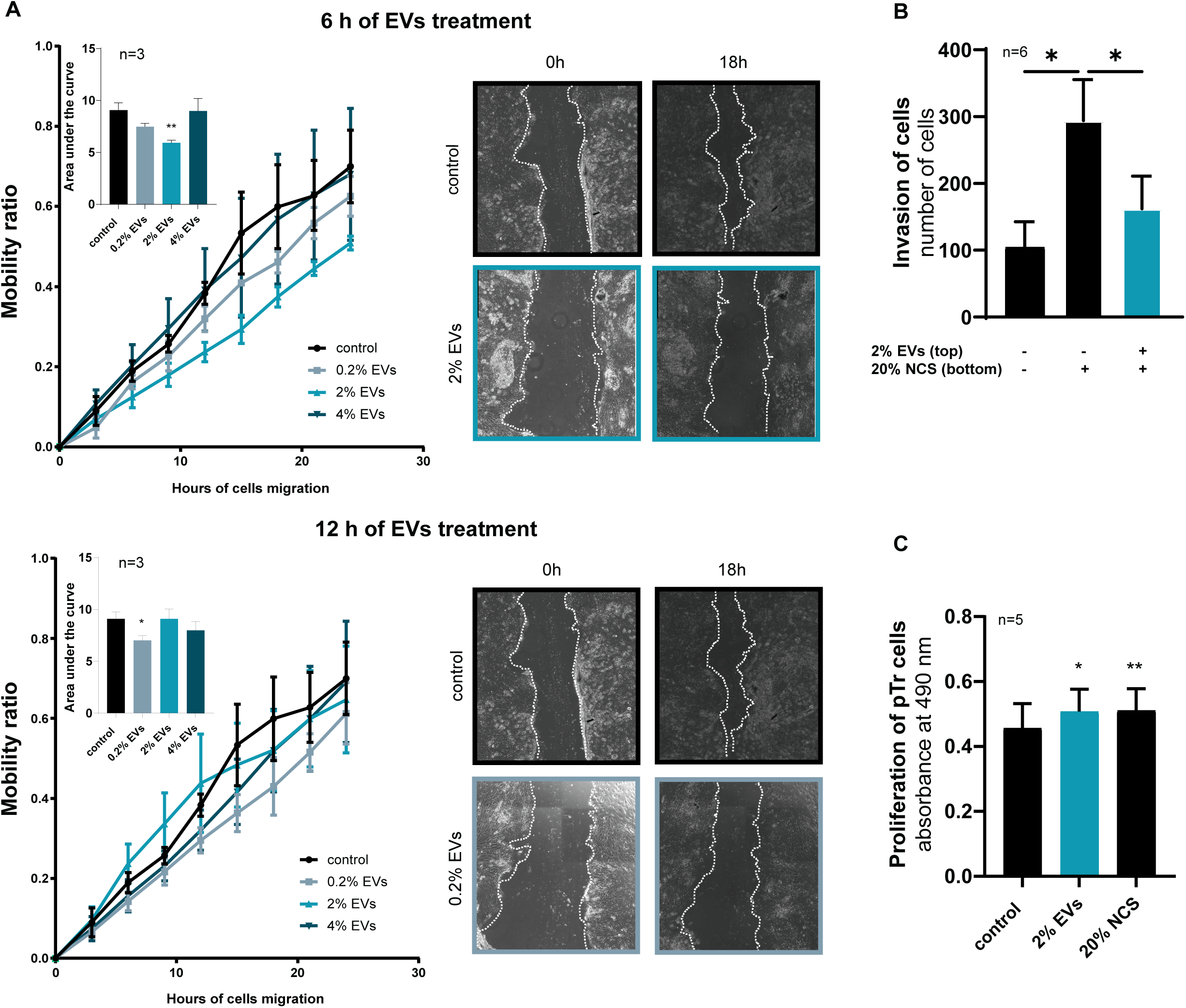
Uterine-derived EVs affect trophoblast cells’ migration, invasion and proliferation. **A**. The migration of pTr cells tested in the wound-healing assay in time and dose-dependent manner. After EVs treatment of pTr cells with three different doses (0.2%, 2%, 4%) for 6 h (top panel) or 12 h (bottom panel), scratch-induced migration was observed for 24 h and snap shots were taken every 3 h. Mobility ratio was calculated at each time point and presented (line graph). Representative images at time 0 and 18 h are shown. The area under the curve for each condition was calculated (column bar graph). * P = 0.0255; ** P = 0.0018 (repeated measures one-way ANOVA and Dunnett’s multiple comparisons test). **B**. The number of pTr cells invaded through Matrigel-coated porous membrane. NCS was used as a chemoattractant. * P ≤ 0.05 (repeated measures one-way ANOVA and Dunnett’s multiple comparisons test). **C**. Proliferation of pTr cells after treatment with 2% of EVs (vs. control). NCS was used as a positive control. * P = 0.0134; ** P = 0.0096 (repeated measures one-way ANOVA and Dunnett’s multiple comparisons test)

### Uterine-derived EVs evoke transcriptomic changes in trophoblast cells related to cell growth, development and interactions

RNA-Seq was performed to get a broader picture of transcriptomic changes in pTr cells evoked by EVs. As a result of EVs’ in vitro autotransplantation to pTr cells, 17 down- and 20 upregulated transcripts were detected (Fig. 3A). Validation of RNA-Seq results using RT-qPCR showed positive correlation for 9 out of 10 genes tested (r = 0.8031, P = 0.0091; YPEL3 was not detected by the assay used in RT-qPCR; Supplementary Fig. 1).

**Figure 3.**
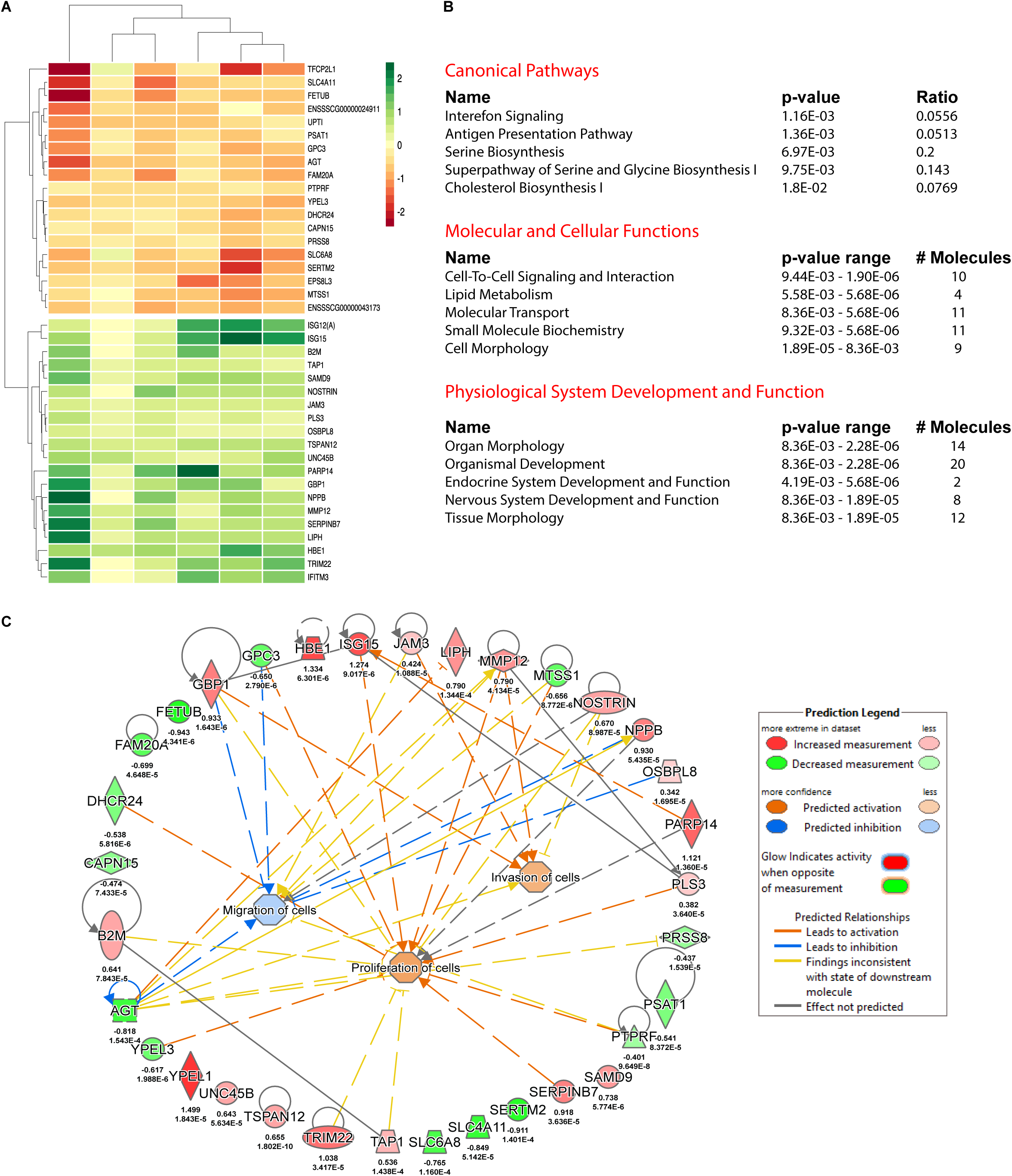
Extracellular vesicles affect the transcriptome of pTr cells. **A**. Heatmap for paired samples clustered according to differentially expressed genes after EVs treatment of pTr cells. Legend presents log2 fold change values (ctrl vs treatment). **B**. Top five canonical pathways (top), molecular and cellular functions (middle) and physiological system development and function (bottom) significantly enriched in pTr cells after EV. Ratio denotes the number of significantly expressed genes compared to the total number of genes associated with the canonical pathway. P values were calculated with a right-tailed Fisher’s Exact Test. **C**. The molecular activation prediction network created using Ingenuity knowledge base and differentially expressed genes in pTr cells after treatment with uterine-derived EVs (vs. ctrl). Colours indicate predicted relationships of gene expression levels and bio-functions, and colour intensities reflect the degree of gene expression or bio-function activity (see prediction legend for details).

The Ingenuity knowledge base was used to identify the canonical pathways and biofunctions enriched by differentially expressed genes (Fig. 3B). Among the top canonical pathways were those important to the immune system, that is, interferon signaling (e.g., ISG15, TAP1; P = 1.16E-03) and antigen presentation (e.g., B2M, TAP1; P = 1.36E-03), as well as in biosynthesis, that is, serine biosynthesis (P = 6.97E-03) and the superpathway of serine and glycine biosynthesis I (e.g., PSAT1; P = 9.75E-03) and cholesterol biosynthesis (e.g., DHCR24; P = 1.8E-02). The top enriched biofunctions were those associated with (i) molecular and cellular functions, such as cell-to-cell signaling and interaction (10 genes; e.g., AGT, GPC3; P range: 9.44E-03 - 1.90E-06); molecular transport (11 genes, e.g., SLC6A8, NPPB; 8.36E-03 - 5.68E-06) and (ii) physiological system development and function, such as organ morphology (14 genes, e.g., FAM20A, SERPINB7; 8.36E-03 - 2.28E-06) and organismal development (20 genes, e.g., MTSS1, ISG15; 8.36E-03 - 2.28E-06). In silico simulation, using the Ingenuity knowledge base and RNA-Seq results, was partially consistent with in vitro autotransplantation assays for principal processes governing conceptus growth, spacing in the uterus, and implantation (Fig. 3C). Predictions highlighted anticipated inhibition of cell migration and stimulation of cell proliferation (more confident) but unexpected stimulation of invasion (less confident).

### Uterine-derived EVs carry unique miRNA cargo important for organismal and embryonic development

Based on our previous studies showing the importance of miRNAs at the embryo–maternal interface (23, 36) and the known fact that miRNAs are transported via EVs, we decided to investigate if, in the physiological context of early pregnancy, these noncoding RNAs can modulate trophoblast cell behavior. First, we analyzed the size distribution of RNA cargo (Fig. 4A). Peaks of the ribosomal RNA were not present, and a clear shift toward shorter RNA fragments was noticed, indicating mostly small RNAs and no cellular RNA contamination in our EV preparations from D12 to D14 and D16 of pregnancy. Neither total RNA nor small RNA concentrations varied among consecutive days of pregnancy. Small RNA fraction consisted of about 30-40% of miRNA, and there was no difference among analyzed days of pregnancy.

**Figure 4.**
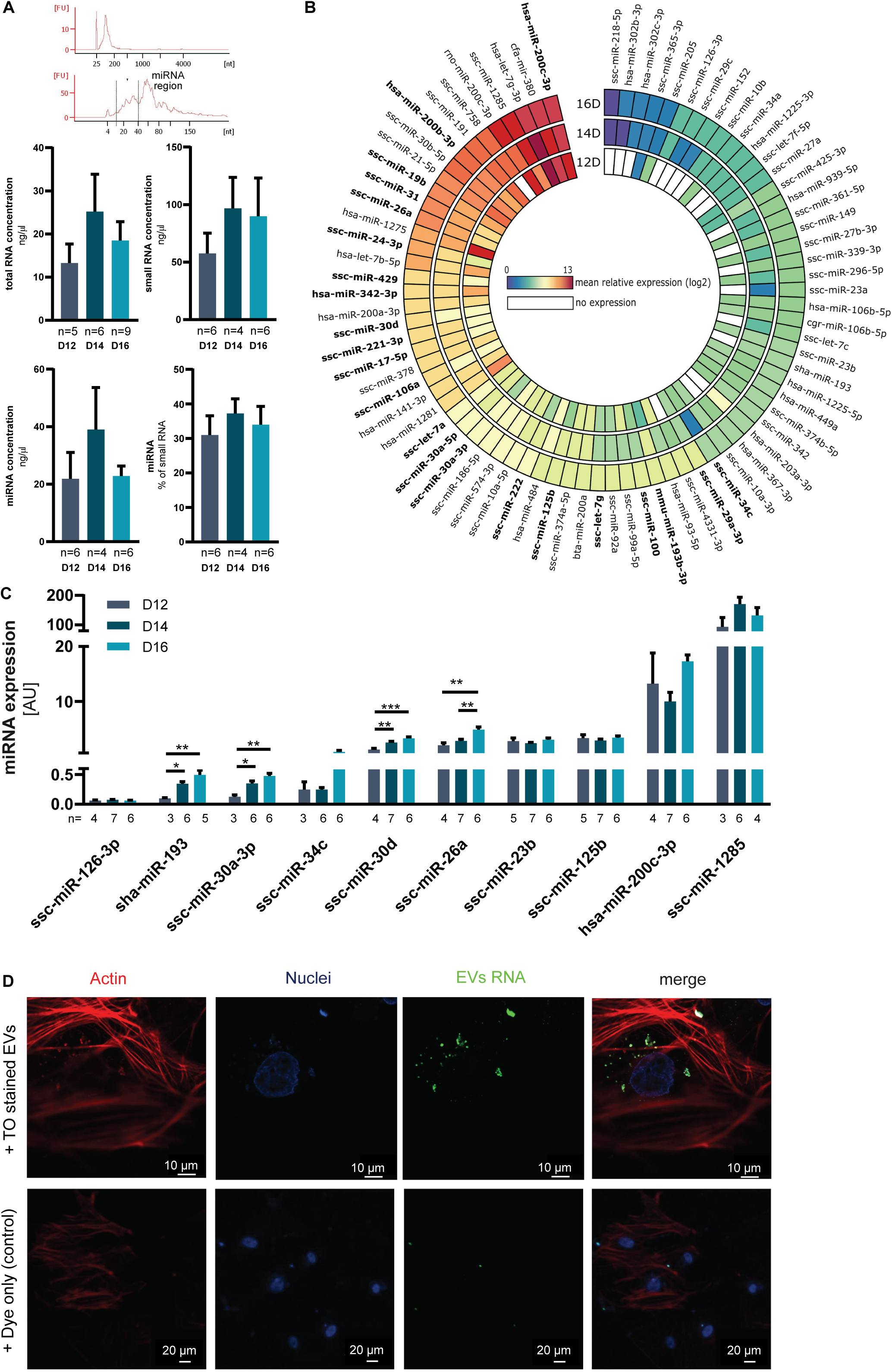
Uterine-derived extracellular vesicles carry miRNA important for organismal and embryonic development. **A**. Representative Bioanalyzer electropherograms of EVs RNA generated using the Agilent RNA 6000 Pico (upper) and small RNA (lower) Kits. Graphs show concentrations of total, small and microRNA as well as % of miRNA in total RNA for EVs collected from uterine lumen during consecutive days (D) of pregnancy (D12, D14 and D16; one-way ANOVA and Tukey’s multiple comparisons test). **B**. Circos plot showing the miRNA abundance in uterine-derived EVs collected on D12, D14, and D16 of pregnancy. The color scale of the heatmap pictures the abundance level as shown in the legend. miRNAs differentially expressed between tested days are marked in bold (one-way ANOVA and Tukey’s multiple comparisons test; detailed results are available in Supplementary Table 2). **C**. Ten miRNAs detected in TLDA analysis were validated using qRT-PCR. * P < 0.05; ** P < 0.01; *** P < 0.001 (one-way ANOVA and Tukey’s multiple comparisons test; detailed results are available in Supplementary Table 2). **D**. Confocal microscopy images showing uptake of labelled EVs’ RNA by pTr cells. RNA cargo carried by EVs was stained with thiazole orange (green). After F-actin was stained with Alexa Fluor 488 Phalloidin (red). Nuclei were stained with Hoechst 33342 (blue).

Next, we screened miRNA cargo in EVs from D12, D14, and D16 of pregnancy using TLDA cards covering miRNAs previously detected in either the endometrium or embryo/trophoblast (23, 36, 57–61). Out of 166 investigated mature miRNAs, 79 were detected in at least two out of three analyzed days of pregnancy (Fig. 4B; Supplementary Table 2). Among them, 22 miRNAs showed differential abundance in uterine-derived EVs in tested days of pregnancy. Additionally, 19 miRNAs were not detected on D12 in contrast with D14 and D16. Abundance of the majority miRNAs increased in the consecutive days of pregnancy. The results of RT-qPCR for selected miRNAs were consistent with TLDA cards (Fig. 4C, Supplementary Table 2).

The Ingenuity knowledge base was used to identify the biofunctions enriched by miRNAs present in EVs on D16. However, 66 out of 79 miRNAs were mapped within the core expression analysis pipeline, as some miRNAs were grouped into families, and some could not be identified due to interspecies differences. Top enriched processes were involved in organismal (20 miRNAs; P range: 3.51E-02 – 2.51E-12) and embryonic development (9 miRNAs; 3.51E-02 – 8.471E-12; Supplementary Table 3).

To test if miRNAs transported via EVs can be taken up by pTr cells after in vitro autotransplantation, nucleic acids cargo was stained with thiazole orange. Indeed, already after 3 h of treatment, stained nucleic acids were visible in cell cytoplasm, whereas for a negative control, any signal was detected (Fig. 4D).

### miRNA–mRNA interactions govern embryo development and function during the peri-implantation period

In the final approach, the physiological relevance of miRNA–mRNA interactions at the embryo–maternal interface was tested in silico using miRNAs detected as EV cargo and transcriptomic datasets available for porcine conceptuses at D12–16 of pregnancy. The list of miRNAs detected in the uterine-derived EVs on D16 was submitted to the miRWalk database to generate a list of potential targets and miRNA-mRNA interactions. Data acquired by Zhang and coworkers (52) were reanalyzed and used to determine genes downregulated in conceptuses on D15 (vs. D12). From the lists of potential EVs miRNA targets, only genes downregulated on D15 conceptuses were used. In total, more than 5000 miRNA–mRNA predicted interactions were identified. The top 5 sub-networks are presented in Figure 5A. Interestingly, among them, there are miR-26a-5p and miR-125b-5p, which we recently identified as important modulators of trophoblast cell function (62).

**Figure 5.**
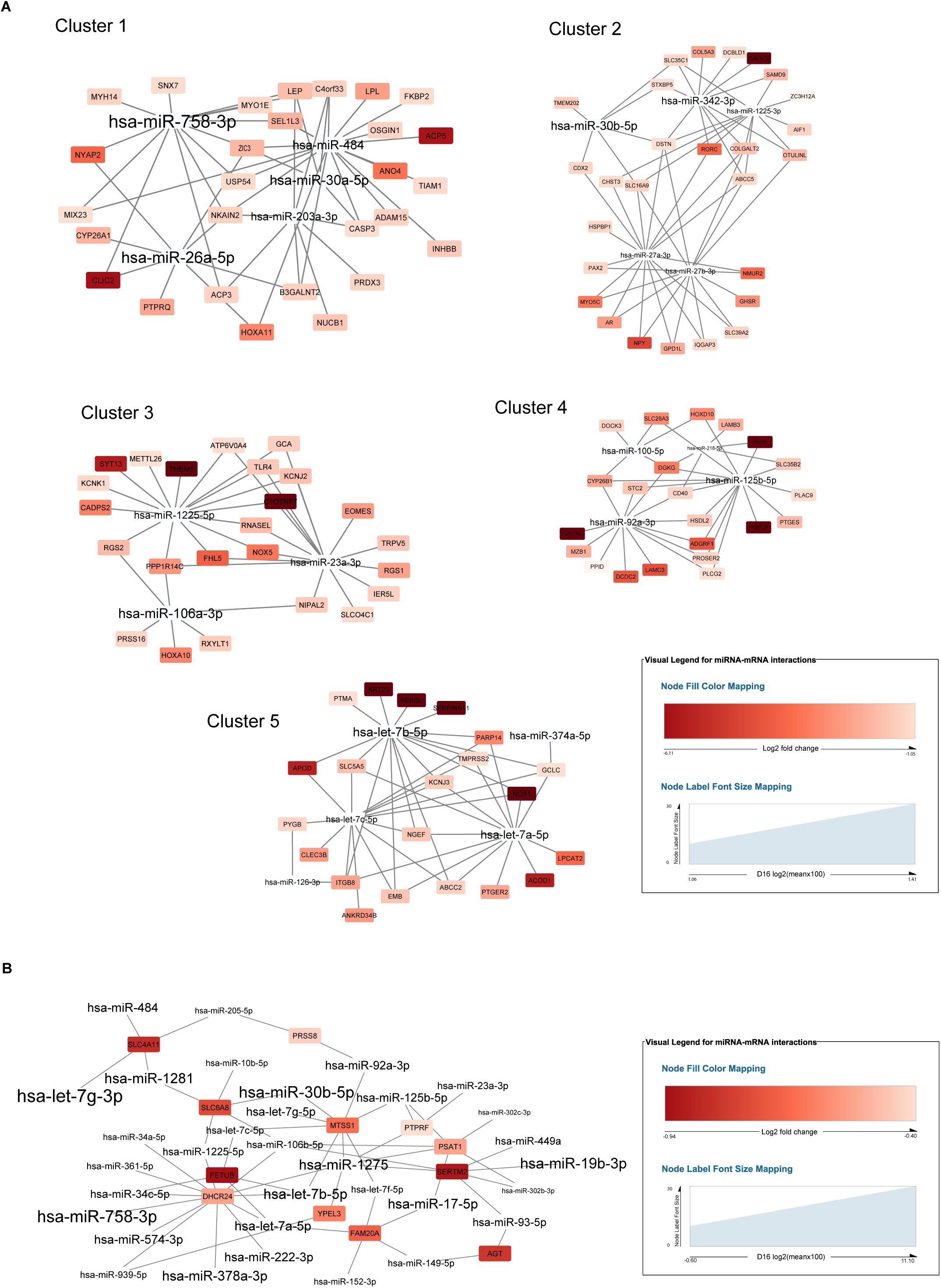
Predicted interaction of miRNAs carried by uterine EVs and mRNAs in conceptuses/trophoblasts at the prei-mplantation stage. **A**. Top 5 subnetworks generated by clustering interactions of miRNAs detected in uterine-derived EVs and genes differentially expressed in conceptuses on day (D) 15 of pregnancy (vs. D12). The miRNA names have different font size based on their abundance (log2 [mean x 100]) at D16 in uterine-derived EVs. Each label is colored based on log2 fold change in gene expression in conceptuses at D15 (vs. D12). **B**. Possible miRNA-mRNA interactions that occurred after pTr cells treatment with EVs. The EVs’ miRNA names have different font size based on their abundance (log2 [mean x 100]) at D16 of pregnancy. Each label is colored based on log2 fold change in gene expression in pTr cells after EVs treatment (vs. ctrl).

Furthermore, we decided to examine possible interactions between miRNAs transported via EVs and genes differentially expressed in pTr cells in an in vitro autotransplantation experiment (Fig. 5B). From the list of potential targets of miRNAs present in EVs at D16, genes showing downregulation in pTr cells after EV treatment were identified. Out of 17 downregulated molecules, 12 were identified as potential targets of miRNAs transported via EVs. Interestingly, among them, miRNAs highly abundant in D16 EVs were observed (i.e., hsa-let-7g-3p, hsa-miR-758-3p, and hsa-miR-30b-5p). Possible interactions were also identified for miRNAs less abundant in D16 EVs, such as hsa-miR-302b-3p, hsa-miR-302c-3p, and hsa-miR-205-5p. Altogether, we showed that miRNAs carried by EVs at the embryo– maternal interface are potent regulators of conceptus transcriptome, affecting embryo development and function.

## Discussion

Growing research on the biological role of EVs provides increasing evidence about their ubiquity. As such, EVs are considered an important element of the embryo–maternal dialog during pregnancy, when reproductive success is determined by the exchange of various molecules. Despite increasing effort, still little is known about EVs’ cargo and their precise mode of action at the peri-implantation phase, characterized by embryo apposition and adhesion common between mammals. Our results demonstrate that EVs containing numerous miRNAs are abundant in the uterus during early stages of pregnancy and that pTr cells respond to molecular messages delivered by EVs via transcriptomic and functional changes critical for pregnancy success.

EVs, as a relatively recent discovery, are rapidly gaining interest and demand that scientific society meets high standards of purification; at the same time, they face numerous methodological obstacles. There is an increasing need for rigorous EV science confirming the existence of a specific and pure population of EVs and defining their role in physiological pathways. Several minimal experimental requirements on how to characterize the EVs (63) are respected in the presented study to authenticate and support our results. The comprehensive characterization of EVs with variety of methods has proven the great quality and purity of samples, allowing us to proceed to further functional and molecular studies on the way to understand EVs’ role at the embryo–maternal interface.

Using our unique model of in vitro autotransplantation, where uterine lumen EVs and trophoblast cells are isolated from the same animal, we were able to observe the physiological response of pTr after EVs’ internalization (i.e., decreased cell migration and invasion, and increased proliferation rate). Interestingly, the morphology of porcine conceptuses changes during the peri-implantation period, when spherical (0.5–1 mm in diameter) blastocysts elongate quickly between D10 and D16 into a 1000-mm filamentous form. These morphological changes are supported by both cellular hypertrophy and hyperplasia (10). Although blastocysts of the pig can retain invasive properties (64), they exhibit noninvasive implantation. Here, we show that uterine lumen EVs are capable of inhibiting pTr cell migration and invasion rate and as such could provide one of the mechanisms crucial in maintenance of pregnancy to term, preventing embryos from unwanted invasion of the uterine wall. Hu and coworkers (65) indicated that spontaneously immortalized porcine trophectoderm cells derived from D12 conceptuses and incubated for a longer time with EVs collected on D15 of pregnancy also exhibited decreased migration. Unfortunately, these observations were not supported by further investigation of possible molecular interactions governing this cellular phenotype. On the other hand, our RNA-Seq analysis revealed 17 down- and 20 upregulated transcripts in primary pTr cells after autotransplantation of EVs to in vitro culture. Interestingly, among the top canonical pathways affected by EVs were those involved in interferon signaling and antigen presentation, which is in agreement with the central role of the immune system at the implantation site in mammals (66, 67). Furthermore, the critical role of amino acids, including serine and glycine (both affected by EV treatment), during rapid conceptus growth and development has been suggested for several species (68– 70). Our in silico simulation of physiological pathways that utilized transcriptomic changes in pTr cells exposed to EVs showed partial consistency with in vitro autotransplantation assays focused on the evaluation of the cellular phenotype. Consistently, stimulated cell proliferation and inhibited cell migration were observed both in silico and in vitro. In contrast, cell invasion inhibited in vitro was not predicted in silico. Physiological cell invasion by definition requires cellular movement (i.e., migration) (71). Unfortunately, IPA prediction failed to show the possibility of simultaneous activation of invasion and inhibited migration of cells, pointing at the strong necessity of in silico prediction validation. Altogether, we showed that EVs at the embryo–maternal interface affect the migration, invasion, and proliferation of trophoblast cells. This is accompanied by changed expression of several genes governing these cellular processes and other pathways known to be essential for proper establishment of early mammalian pregnancy.

EVs carry a wide range of RNAs representing many biotypes (72). The major fraction carried by EVs isolated from the uterine cavity during the peri-implantation period were small RNA species. Further analysis of small RNAs showed that 30–40% is covered by miRNAs. TLDA cards allowed us, with high sensitivity and specificity, to quantify mature miRNA EV cargo of either endometrial or embryo/trophoblast origin, previously detected by us and others (23, 36, 57–61). Among 116 miRNAs assayed, 79 were detected in at least two out of three analyzed days of pregnancy. Interestingly, 19 miRNAs were absent on D12, whereas their expression was detected on D14 and D16. This pattern of expression also proves the intense molecular changes occurring between D12 and D16 at the embryo–maternal interface when the embryo elongates and the endometrium remodels to reach the full capacity to accept the implanting embryo. Importantly, miRNAs detected as EV cargo were identified as involved in organismal and embryonic development, which we further validated at the cellular level using a unique model of in vitro autotransplantation, allowing us to track the consequences of EV delivery to pTr cells within the same animal.

Identification of the possible miRNA–mRNA interactions based on ex vivo EVs’ miRNA abundance and conceptus transcriptome data at the peri-implantation period (52) showed thousands of possible interactions. This agrees with the known fact that a particular miRNA can target many different mRNAs (73), and particular messenger RNA can bind to a variety of miRNAs, either simultaneously or in a context-dependent fashion (74, 75). Knowing these limitations in data interpretation, still considering an existing coexpression of the miRNA and its targets is one of the recommended strategies for possible interaction identification (76). Fortunately, among top miRNA–mRNA networks, we were able to find two miRNAs, miR-26a-5p and miR-125b-5p, suggested by our previous studies, to be important players in early pregnancy events (23, 37, 38). Our further investigation proved recently that both miRNAs are essential modulators of trophoblast cell function (62). Interestingly, one network was composed of miRNAs (hsa-let-7a-5p, hsa-let-7b-5p, hsa-let-7c-5p) belonging to the let-7 family, highly conserved across species in sequence and function (77). Among genes targeted by these miRNAs are those the most downregulated in conceptuses at D15 (vs. D12), such as *KRT25* (Keratin 25), *RERGL* (Ras-Related and Estrogen-Regulated Growth Inhibitor-Like Protein), *SERPINB11* (Serpin Family B Member 11), and *NOS1* (Nitric Oxide Synthase 1). Here, we show that miRNAs showing both high and low abundance in EVs are potent regulators of gene expression at the site of embryo–maternal crosstalk. As stressed before, miRNA–mRNA interactions are context dependent, and numerous factors, such as the number of miRNAs simultaneously targeting one target, can influence the interplay. Thus, we developed the autotransplantational model used in this study to employ a unique composition of miRNAs present in the uterine environment on D16 of pregnancy and observe widespread transcriptomic and functional changes in trophoblast cells of the same animal, giving us the chance to make observations as close to the naïve state as possible.

The presented data support the idea of the crucial role of EVs in a complex and multidimensional embryo-maternal communication occurring during early pregnancy. We proved that EVs transport a wide range of miRNAs in the consecutive days of the peri-implantation period, with the potential to affect physiological pathways. We showed that these EVs are important players in governing embryo growth and development, as well as early stages of implantation. Thus, we conclude that the unique EV population present in the uterine cavity during pregnancy is the key to the success of implantation and pregnancy.

## Acknowledgements

The authors are grateful to Dr A. Nitkiewicz, Dr K. Witek, M. Guzewska, K. Drzewiecka, M. Romaniewicz, P. Golder, M. Sikora, and K. Gromadzka-Hliwa from the Institute of Animal Reproduction and Food Research, Polish Academy of Science for their excellent technical assistance laboratory; Dr E. Karnas, from the Laboratory of Stem Cell Biotechnology, Malopolska Centre of Biotechnology, Jagiellonian University for performing Nanoparticle Tracking Analyses; M. Angenitzki from The Hebrew University of Jerusalem for invaluable help with electron microscopy imaging.

## Author Contributions

J.S. designed and performed experiments, collected, analyzed, and interpreted data; drafted the manuscript and participated in the preparation of its final version. J.S. K.M. and M.M.K contributed to the bioinformatics analyses. Z.P.R designed TLDA cards and established protocol for EVs isolation. Y.H. was responsible and supervised the transmission electron microscopy imaging, and participated in the preparation of final version of manuscript. M.M.K. conceived and supervised the study, designed experiments, analyzed, and interpreted data and was responsible for the final version of the manuscript.

## Additional Information

The authors declare no competing or financial interests. Supplementary information is available for this paper.

